# Reintroducing genetic diversity in populations from cryopreserved material: The case of Abondance, a French local dairy cattle breed

**DOI:** 10.1101/2023.01.17.524356

**Authors:** Alicia Jacques, Grégoire Leroy, Xavier Rognon, Etienne Verrier, Michèle Tixier-Boichard, Gwendal Restoux

## Abstract

**Background:** Genetic diversity is a necessary condition for populations to evolve under adaptation, selection or both. However, genetic diversity is often threatened, in particular in domestic animal populations where artificial selection, genetic drift and inbreeding are strong. In this context, cryopreserved genetic resources appear as a promising option to reintroduce lost variants and to limit inbreeding. However, while more common in plant breeding, the use of cryopreserved resources is less documented in animals due to a longer generation interval making it difficult to fill the performance gap due to continuous selection. Thus, this study investigates one of the only concrete cases presenting the results of the use of cryopreserved semen of a bull born in 1977 and belonging to a disappeared lineage, into the breeding scheme of a French local dairy cattle breed, the Abondance breed, more than 20 years later.

**Results:** We found that this re-used bull was very original relative to the current population and thus allowed to restore genetic diversity lost over time. The expected negative gap in milk production due to continuous selection has been absorbed in a few years by preferential mating with elite cows. Moreover, the re-use of this old bull did not increase the level of inbreeding, it even tended to reduce it by avoiding mating with relatives. Finally, the reintroduction of an old bull in the breeding scheme allowed for increased performance for reproductive abilities, a trait that was less subject to selection in the past.

**Conclusions:** The use of cryopreserved material was efficient to manage the genetic diversity of an animal population, by mitigating the effects of both inbreeding and strong selection. However, attention should be paid toward mating in order to limit the disadvantages associated with the provision of genetic originality, notably a discrepancy in the breeding values for selected traits or an increase in inbreeding. Therefore, a careful characterization of the genetic resources available in cryobanks could help to ensure the sustainable management of populations and of local or small ones in particular. These results could also be transferred to the conservation of wild populations.

## Background

The genetic variability of populations is important for adaptation to changing environments. Indeed, highly diverse populations have a higher probability to contain advantageous or adaptive allelic variations leading to a significantly higher evolutionary potential [1]. In addition, genetic diversity is essential for animal populations under selection since it affects the genetic variance needed to carry out selection. In dairy cattle, this selection is based on the use of a limited number of sires, resulting in both genetic drift and increased inbreeding all over the genome [2]. Thus, the selection process inevitably leads to a more or less rapid reduction in genetic diversity depending on selection intensity and breeding goals [3,4]. Depending on the breed and its breeding scheme, the implementation of genomic selection could have impacted genetic gain as well as genetic diversity, calling thus for a careful monitoring in order to ensure the sustainability of these programs [5–7]. Indeed, monitoring the level of genetic diversity of local breeds is needed to take action based on their risk status [4,8].

Cryopreservation and the development of gene banks, are useful contributions to the conservation of genetic diversity in domestic animals. However, the use of *ex situ* genetic resource conservation programs should be combined with *in situ* breed conservation, as mentioned in the FAO Global Plan of Action [9]. In France, the National Gene Bank was created in 1999 for the conservation of semen and embryos of domestic animal breeds with the aim of hosting samples representative of the genetic diversity of all French breeds. Then, the genetic collections rapidly grew, with a major contribution of the bovine species [10]. One of the main purpose of cryopreservation is the conservation of genetic diversity of threatened populations only as backup, with one indicator of the Sustainable Development Goals (SDG 2.5.1b) considering the number of breeds with an amount of genetic material stored sufficient to reconstitute the breed in case of extinction [11]. Cryopreserved material can also be used for other goals such as the reintroduction of genetic diversity to limit inbreeding in targeted populations [12]. However, the effective use of these resources for that purpose has remained quite low, although a concrete test already demonstrated its feasibility in poultry [13].

The long term cryoconservation of reproductive material (semen, ova, embryos) makes possible to use ancient individuals in order to re-introduce genetic diversity in on-farm populations [14]. However, the use of ancient genetic resources can hinder genetic progress for traits that are currently being selected [15]. The more a population has been subjected to strong selection, resulting in large genetic gain over successive generations, the more the conserved genetic resource will exhibit a lag in performance for the selected traits. While it is more common to use external genetic resources in current plant breeding program, it is almost never the case for animals where the longer generation interval reduces the efficiency of bridging or pre-breeding strategies to fill the performance gap [16,17]. However, a theoretical study showed that the use of old Dutch cryobank bulls could increase genetic variability and still improve genetic merit in a current population of Holstein Friesian dairy cattle [18]. Another risk of re-using ancient sires for reproduction is the increase of the inbreeding level of the populations by mating them with some of their relatives in the current population. A real compromise has to be found between a contribution to genetic diversity on the one hand, and the slowing down of genetic progress and the impact on inbreeding on the other hand. Simulation studies have already shown the potential of using cryopreserved genetic material to bring back diversity [19] but very few real cases have been reported up to now [20]. A recent case illustrated the successful recovery of a lost lineage of the Y chromosome in the Holstein breed thanks to the use of frozen semen from the National Gene Bank in the USA [21].

In this study, we propose to analyze a concrete case of the use of an ancient cryopreserved bull to restore genetic diversity in a selected French local dairy cattle breed, the Abondance breed. We used pedigrees, genotypes and genomic estimated breeding values to determine the impact of the re-use of this bull on the current Abondance breeding scheme from both a diversity and a performance perspective. The objective was to prove the effectiveness of using *ex situ* genetic resources to reintroduce genetic diversity and to understand the key parameters related to a successful use of cryopreserved collections.

## Methods

### Description of breed and animals

The Abondance breed originates from the Chablais region, which is the northernmost part of the French Alps, located between the French shoreline of Lake Geneva and the Valais canton in Switzerland. The geographical isolation of this area has favored the development of an original and hardy cattle population adapted to the mountain environment and to farming systems involving transhumance (see Additional file 1, Figure S1). About 90% of its milk is processed in cheese, mainly cheeses benefiting form a Protected Designation of Origin (PDO). The Abondance breed is the fourth French dairy cattle breed in population size, with about 48,000 cows, but representing only 1.4% of the total French dairy herd. Selection though progeny testing has been conducted for decades and thus, pedigrees and performances are available for many individuals. Genomic selection is now applied since 2015 giving access to genomic estimated breeding values (GEBV) comparable between animals of different cohorts. Finally, cryopreserved material is available in collections in breeding companies and at the French National cryobank. Crossbreeding with Red Holstein bulls took place during the 1980s in order to improve both milk yield and udder morphology. Then, some crossbred AI bulls (75% Abondance, 25% Red Holstein) were available from the end of the 1980s to mate cows on farm. Conversely, some purebred families no longer gave approved sires, such as the Amiens bull lineage. Finally, this strategy was discontinued at the end of the 1990s and a forced return to a pure breed may have decrease the genetic variability of the breed due to a bottleneck effect. This trend motivated the breeders’ association to reintroduce genetic diversity by the use of cryopreserved material at the beginning of the 2000’s. The Abondance breed was therefore a good case study to investigate the consequences of reintroducing genetic diversity through the use of former breeding stock.

In the present study, we focused on the use of Naif semen, a bull born in 1977 and a son of Amiens, whose semen was cryopreserved. Its semen has been used in two distinct periods, first between 1980 and 1993 and then between 2004 and 2009. We defined two cohorts of contemporary genotyped sires. Cohort 1 corresponds to 62 sires, born between 1970 and 1991, which produced offspring between 1980 and 1993 along with Naif. Cohort 2 corresponds to 165 sires, born between 1982 and 2007, that produced offspring between 2004 and 2009, corresponding to the period when Naif was re-used. Then, we defined four female cohorts corresponding to the dams which produced progeny during the same two periods and for which the values of two indices (the dairy merit index and the milk production) were available. Cohort 1a corresponds to 2443 dams, mated to sires other than Naif, that produced offspring between 1980 and 1993. Cohort 1b consists in the 37 females mated to Naif during its first period of use. Three females were mated with Naif as well as with another sire for the period 1980-1993 and were thus present in both cohorts. Cohort 2a corresponds to 4092 dams, mated to sires other than Naif, which produced offspring between 2004 and 2009. Cohort 2b is made up of 25 females mated to Naif when its semen was re-used. Fifteen females were mated with Naif as well as with another sire for the period 2004-2009 and were thus present in both cohorts. Finally, we also defined the cohort ‘2017’ as the current population consisting of all individuals born in 2017, in order to study the long-term effect of Naif re-use.

### Pedigree data

We used a dataset that included all the ancestors of available genotyped individuals in 2017 in the Abondance breeding scheme extracted from the national database. Thus, the pedigree included a total of 25,010 individuals born from 1944 to 2018. The quality of the pedigree was evaluated through the equivalent number of generation with the NGEN module of the PEDIG software [22]. Pedigree quality was computed as the average between males and females for the two Naif production periods (1980-1993 and 2004-2009), the two periods corresponding to the birth years of sires from cohort 1 (1970-1991) and cohort 2 (1982-2007), as well as for cohort 2017.

Naif genetic contribution to the gene pool of a given cohort was defined as the probability that, at any neutral locus, an allele drawn at random in the genotype of an animal drawn at random in this cohort originates from Naif. This probability was computed from pedigree data using the PEDIG software. Total Naif contributions were calculated for each year from 1980 to 2017. Then two types of contributions were defined, an old contribution from the first use of Naif (1980-1993) and a recent contribution from the contemporary use of the cryopreserved Naif’s semen (2004-2009). These contributions were calculated using the same method as for the total contributions. The contemporary use of Naif was computed by affecting a new identifier to Naif when used during the second period with both same parents.

### Molecular data

We had the genotypes of 6958 individuals with the 50K SNPs chip (Illumina Infinium^®^ BovineSNP50 BeadChip). Quality control was performed by removing SNPs with a call rate inferior to 99%, as well as individuals which had less than 99% genotyped SNPs. After quality control, no SNPs were deleted and 43,801 autosomal markers remained in the data sets. However, 26 pairs of markers were found to have identical positions in the genome. These 52 SNPs were removed from the analyses. The density of markers averaged to one SNP every 57.2 ± 60.0 kb.

#### Measurements of heterozygosity

The individual observed heterozygosity of Naif, and of sires of the two cohorts were computed and the means heterozygosity of cohort 1 and 2 were compared using a two-way ANOVA.

#### Measurement of inbreeding

Inbreeding was assessed from molecular data using Runs of Homozygosity (ROH hereafter). ROHs represent long autozygous segments of the genome (i.e. identical by descent). Here, a ROH was defined as a homozygous segment of at least 15 SNPs and 1000 kb in length, with at least one SNP every 70 kb. Two consecutive SNPs could not be included in the same ROH if they were separated by more than 140 kb. ROHs were detected using the “homozyg” PLINK 1.9 function [23,24]. The size of the sliding window has been set to 15 SNPs. The number of heterozygous calls in the sliding window was limited to 1, and the limit for missing data was 5. Inbreeding estimates based on the ROH, *FROH_i_*, were calculated according to McQuillan *et al* [25], as the proportion of the genome included in the ROH as follows:

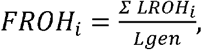

with *Σ LROH_i_* the total length of ROH for individual i and Lgen the size between the first and last marker covering the considered genome.

The 1148 genotyped individuals born in 2017 were grouped according to their relationship with Naif, forming two groups: one group with a recent link to Naif, where it appears on the pedigree to be a father on his second use, and the other group with no link to the recent use of Naif.

#### Genetic Structure by Multivariate Analysis

Principal component analyses, PCA, were conducted with cohorts 1 and 2 using the ade4 package [26]. Naif was added as a supplementary individual to the PCAs (i.e. it is not contributing to the construction of the principal components).

A between-class analysis was then carried out on the 2017 cohort specifying the family links of the individuals with Naif. This analysis aimed to maximize the between groups variance while minimizing within groups variance. The 1148 individuals were separated into 6 classes representing the possible combinations of the different uses of Naif, either the absence of a link, or the presence of one or two old or recent links (consequence of its first or second use through a single parent or both of them). 85 individuals had no family link to Naif (0_LWN), 49 individuals had a recent link to Naif through one of their parents (1_LWN_R), 387 individuals had an old link to Naif through one of their parents (1_LWN_O), 479 individuals were related to the old use of Naif by both parent (2_LWN_O), 1 individual was related to the recent use of Naif by both parent (2_LWN_R) and 147 individuals had one old and one recent link with Naif (2_LWN_OR) (see Additional file 2, Figure S2).

### Genomic performance data evaluated in 2017

Genomic estimated breeding values, GEBVs, were assessed in 2017 for all genotyped individuals. We extracted these values for the cohort 1, cohort 2 and Naif. Genomic performance data are missing for some bulls from cohort 1. Thus, the GEBV calculated for cohort 1 correspond to the 54 bulls evaluated out of the 62 ones available. We also extracted these values for the 15 sons and 25 grandsons of Naif from its second use, as well as for the 155 bulls of the cohort 2 with no genetic link to Naif’s second use, their 416 sons and 528 grandsons for which GEBVs was available.

For male cohorts, we focused on three different multiple-trait indices: (i) the total merit index (ISU, *Indice de Synthèse Unique*), (ii) the dairy merit index (INEL, *Indice National Economique Laitier*) and (iii) the reproduction merit index (REPRO). For female cohorts, we focused on two milk indices, the milk yield (MILK) and the dairy merit index (INEL).

### Statistical analysis

All statistical analyses and graphical representations were made using R [27] and the ggplot2 package [28]. Statistical tests were made using the lm function and post-hoc and comparisons were made using the emmeans package [29], type II ANOVA were performed using the car package [30].

## Results

### Pedigree data

#### Quality of genealogy

The pedigree quality for the two Naif production periods varied from 3.19 (sd=0.41) equivalent number of known generations for the cohort 1980-1993, to 5.80 (sd=0.40) for the cohort 2004-2009. The equivalent number of known generations was 7.57 (sd=0.08) for the cohort of 2017. The equivalent number of known generations for the cohort 1 (1970-1991) and cohort 2 bulls (1982-2007) were 2.96 (sd=0.73) and 4.12 (sd=0.92) respectively.

#### Progeny of Naif

During its first use, Naif produced 45 direct offspring born between 1980 and 1993 (Figure 1). His semen was then used to produce 33 offspring born from 2004 to 2009. Thus, Naif produced 3.2 offspring per year during its first period of use and 5.5 offspring per year when its semen was re-used. He was not used anymore for artificial insemination after 2009.

**Figure 1.**
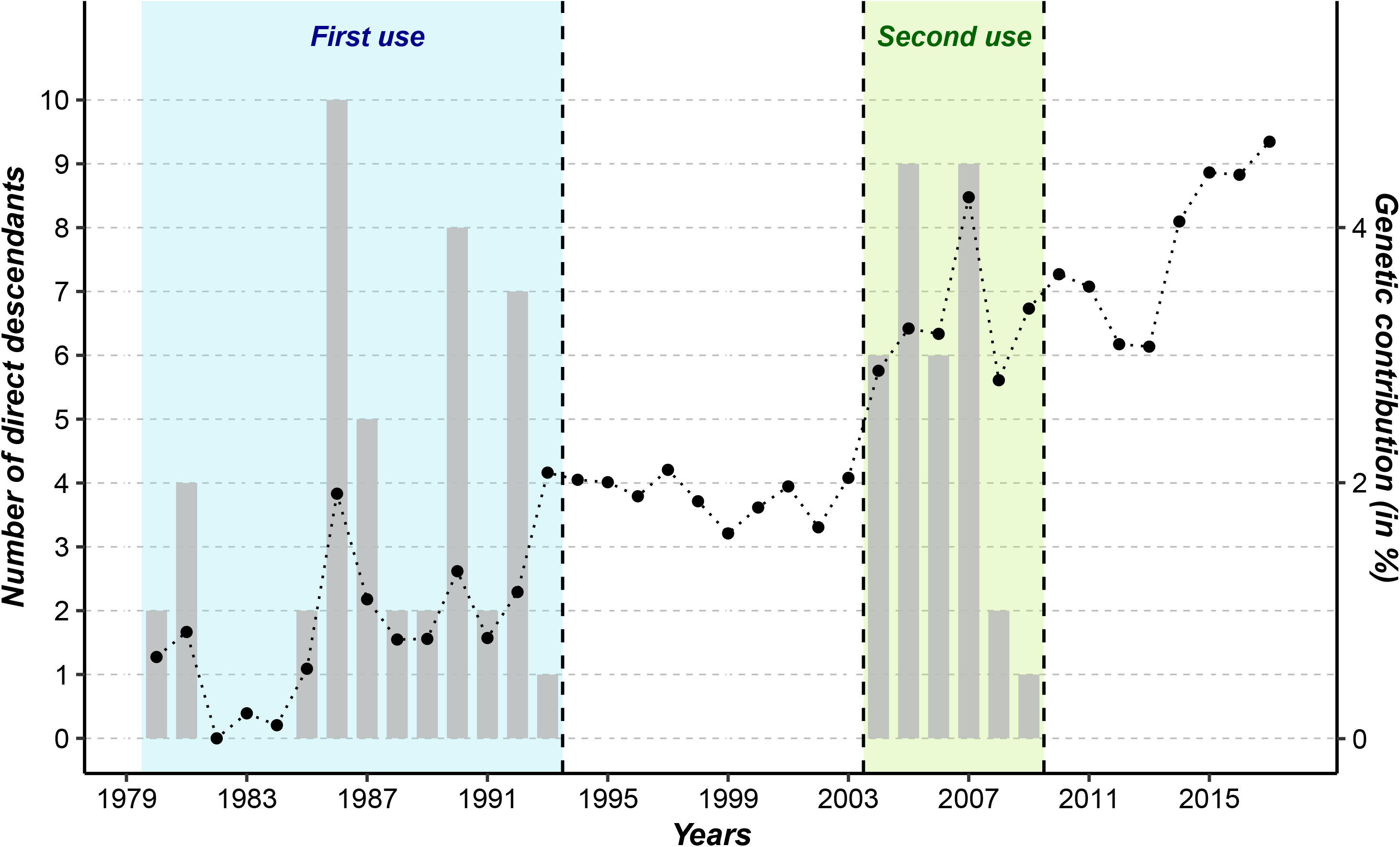
Naif direct progeny production and total annual pedigree-based contribution from 1980 to 2017. Blue: first period of Naif use – Green: second period of Naif use

#### Genetic contribution of Naif

In its first period of use, the overall genetic contribution of Naif increased from 1980 to 1993. From 1994 to 2003, its genetic contribution remained fairly constant. During its second use, Naif’s contribution increased again. From 2009 onwards, a slight decline appeared over the next four years, followed by a marked increase from 2014 to 2017 due to the use of Naif’s progeny (Figure 1). Distinguishing between past and recent Naif contributions (Figure 2) revealed the impacts of the two periods of Naif use: older contributions had a larger impact than recent ones, with the exception of the year 2017 where the recent contribution becomes greater than the old contribution. The recent contribution globally increased over time, in particular from 2013.

**Figure 2.**
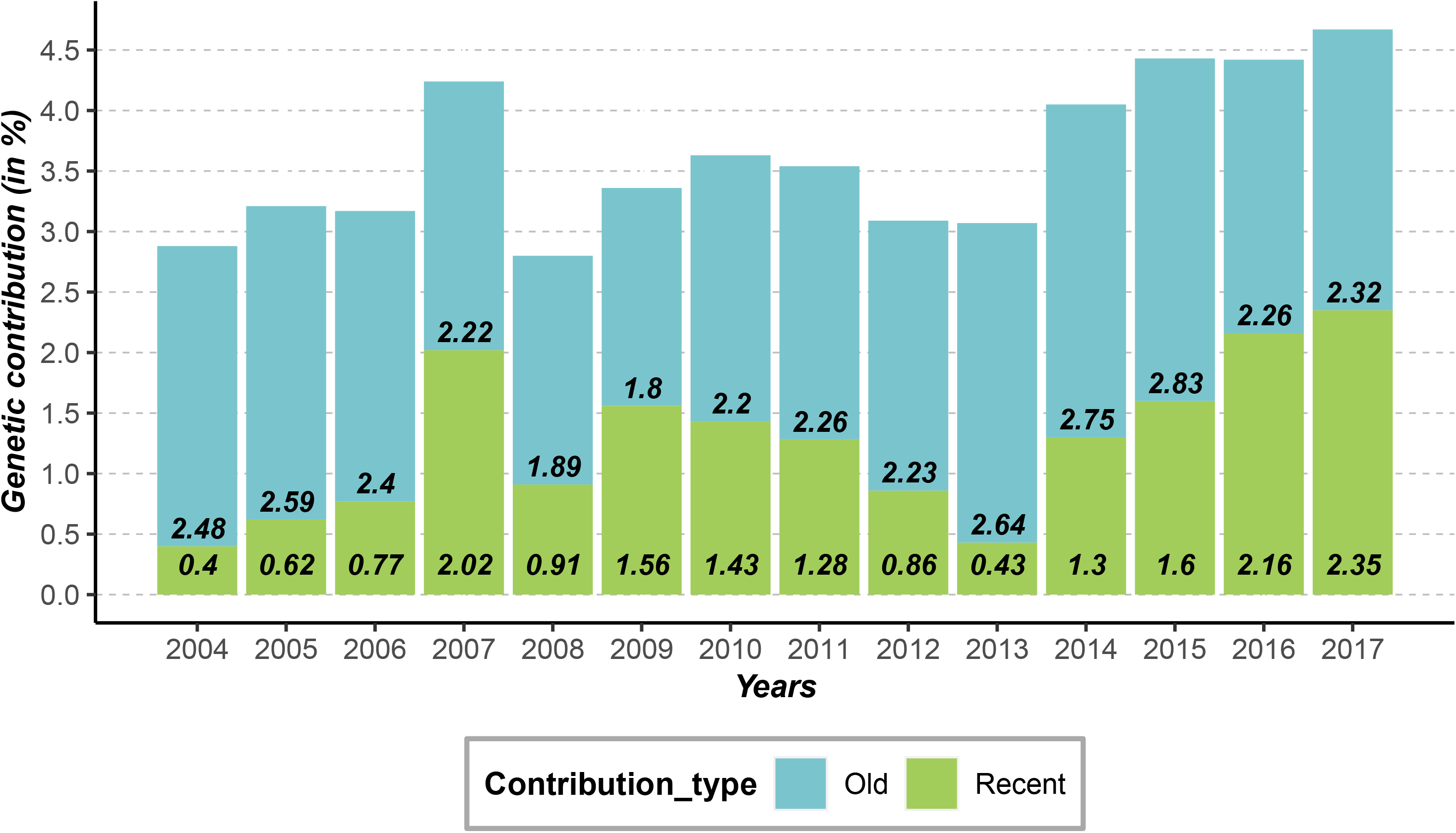
Old and recent contribution of Naif evaluated from pedigree from 2004 to 2017. In blue: old contribution from the first period of Naif use - In green: recent contribution from the second period of Naif use

### Molecular data

#### Measurements of heterozygosity

The average heterozygosity along the genome of the 62 sires from cohort 1 was 32.9% (sd=1.6) while that of the 165 sires from cohort 2 was 31.3% (sd=1.3). Over the whole genome, the average heterozygosity decreased significantly from the first to the second cohort (F=60.5, df=1, p<0.05). Naif has an individual heterozygosity rate of 33.6%, which belongs to the third quartile of the distribution of cohort 1, while he was one of the most heterozygous individuals of the cohort 2 (Figure 3).

**Figure 3.**
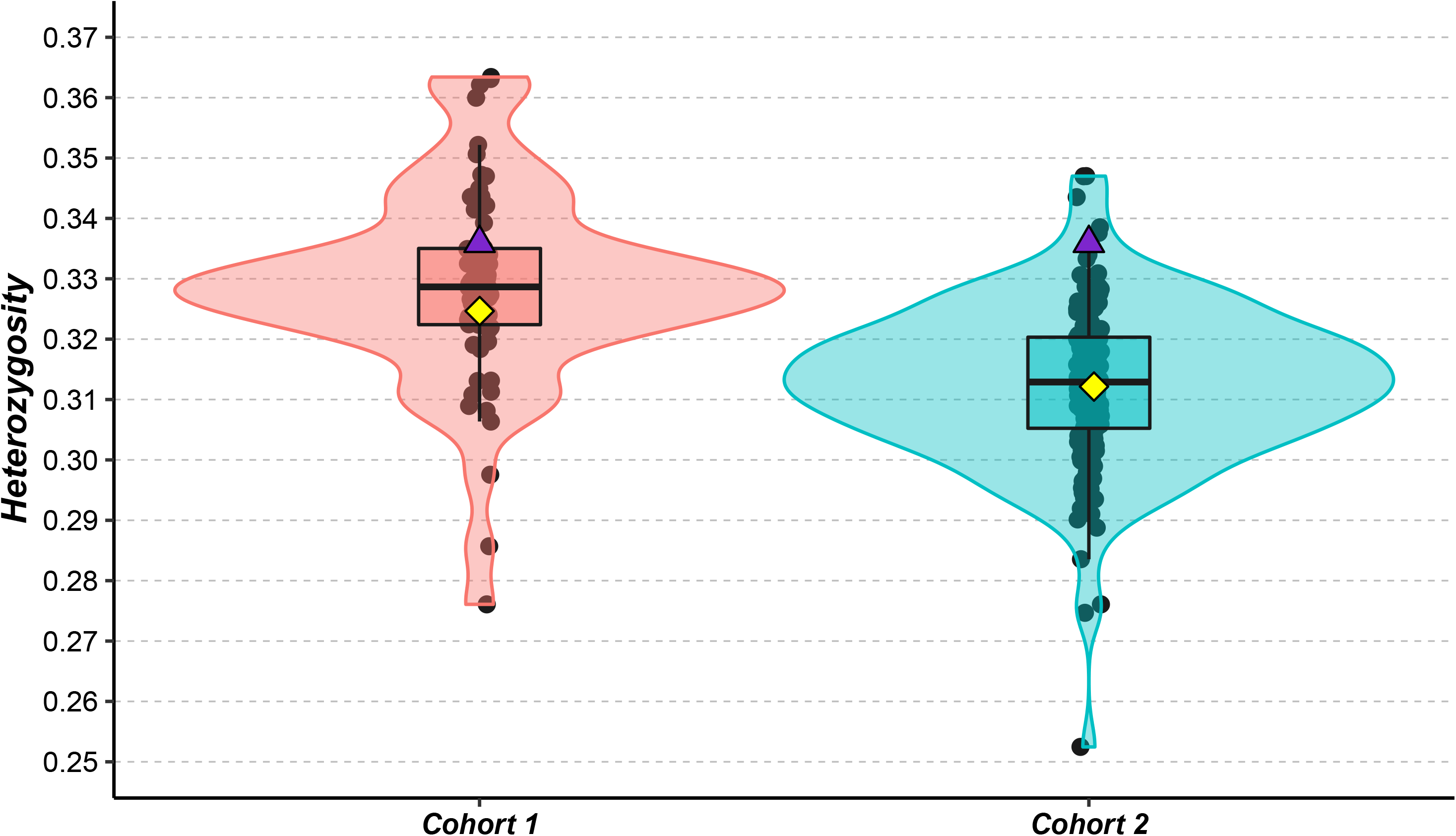
Average heterozygosity of contemporary cohorts at both uses of the Naif bull. The 62 bulls in Cohort 1 are shown in pink, the 165 bulls in Cohort 2 are shown in blue, Naif is represented by the purple triangle, the mean of each cohort corresponds to the yellow square

#### Inbreeding measurements

For cohort 2017, 85 animals were genetically unrelated to Naif (0_LWN), 436 animals had a single family link with Naif through either the maternal or paternal way (1_LWN) and 627 animals had two family links with Naif through both parents (2_LWN), with average inbreeding of 8.67% (sd=1.81), 8.64% (sd=1.58) and 8.58% (sd=1.62) respectively. Inbreeding was not significantly different according to the number of links with Naif (F=0.274, df=2, p=0.76). In addition, 197 individuals had a link to Naif from its second period of use while 951 individuals were not related to the recent use of Naif. Mean inbreeding was 8.74% (sd=1.62) and 8.01% (sd=1.51), respectively for individuals with no link to the recent use of Naif and those related to recent Naif use (Figure 4). A significant decrease in inbreeding was observed following the use of the cryopreserved bull (F=34.06, df=1, p<0.05).

**Figure 4.**
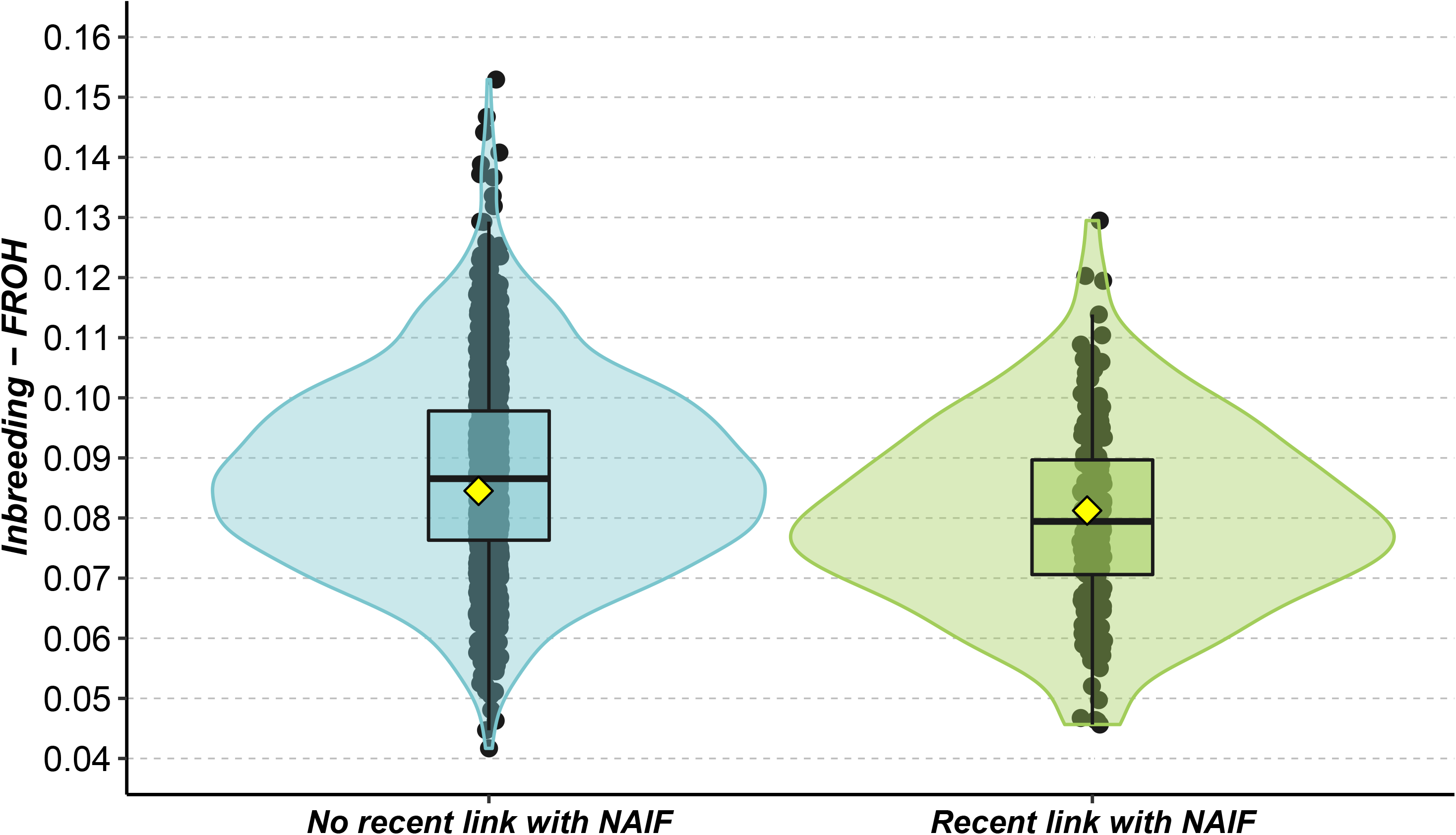
Inbreeding of individuals of cohort 2017 depending on their link with the recent use of Naif. Green: individuals not from recent use of Naif - Blue: individuals from recent use of Naif

#### Genetic Structure by Multivariate Analysis

The two first components of the PCA of cohort 1 explained 11.6% of the total inertia. Naif appears to be an “average” individual within the cohort 1 very close to the origin of the PCA (Figure 5a). Naif appeared more extreme and different from the “average” sires of cohort 2 in the PCA related to its second use (Figure 5b). The two first axes accounted for 6.7% of the total variability. In addition, individuals with a link to Naif (LWN, n=85), i.e. for which Naif is in their pedigrees, appeared separated from individuals with no link with Naif (no_LWN, n=80) on this PCA (see Additional file 3, Figure S3).

**Figure 5.**
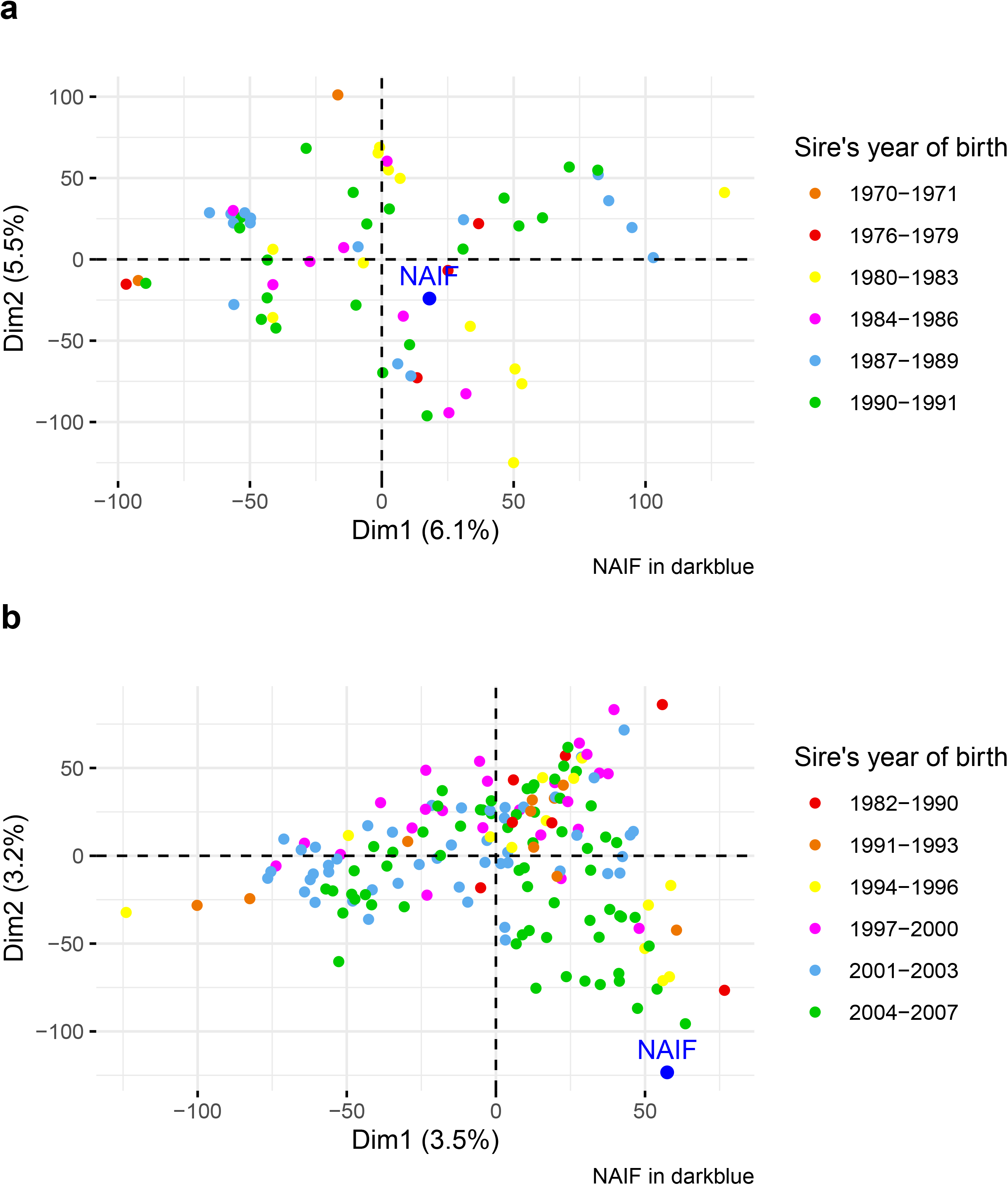
Principal Component Analysis of genotyping data for Cohort 1 (a) and Cohort 2 (b). Naif is represented by the blue dot

For the between-class analysis of the 2017 cohort, the 3 groups with recent use of Naif (1_LWN_R, 2_LWN_R, 2_LWN_OR) appeared separated from the other 3 groups involving no link to Naif (0_LWN) or only links due to its first use (1_LWN_O, 2_LWN_O) (Figure 6).

**Figure 6.**
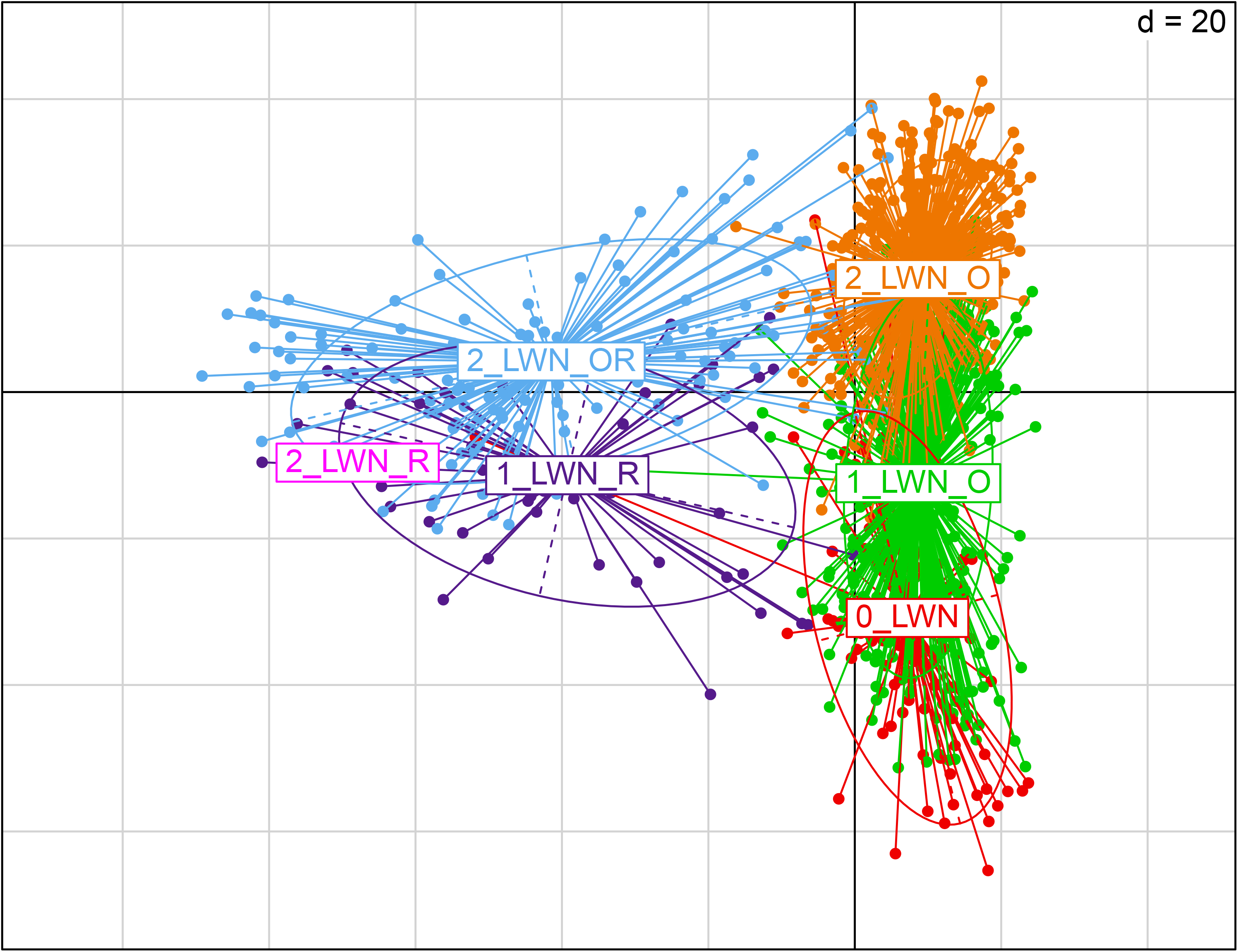
Between-Class Analysis of genotyping data of the cohort 2017. Red: individuals not related to Naif - Green: individuals with 1 old link with Naif - Orange: individuals with 2 old links with Naif - Blue: individuals with 1 old and 1 recent link with Naif - Purple: individuals with 1 recent link with Naif - Magenta: individuals with 2 recent links with Naif

### Breeding values

#### Males

The ISU, INEL and REPRO values of Naif were 70, −24 and 0.8 respectively. The values were 75.46 (sd=15.72), −19.70 (sd=15.85) and 0.26 (sd=0.36) for cohort 1 and 90.75 (sd=16.29), −6.84 (sd=15.86) and 0.13 (sd=0.57) for cohort 2 for ISU, INEL and REPRO respectively. The distribution of these values for cohort 1 and cohort 2 and the relative position of Naif are shown on Figure 7. The means of the ISU and INEL for cohorts 1 and 2 were significantly different (F=36.42, df=1, p<0.05 for ISU, F=26.78, df=1, p<0.05 for INEL). We calculated the difference between the mean of cohort 1 (or cohort 2) and the Naif value for ISU. For ISU, the gap with the cohort 1 was 5.46 and the gap with the cohort 2 was 20.75. For INEL, the gap was 4.30 and 17.16 with the cohort 1 and the cohort 2 respectively. There was no significant difference between the means of the REPRO index for the two cohorts (F= 2.457, df=1, p=0.12).

**Figure 7.**
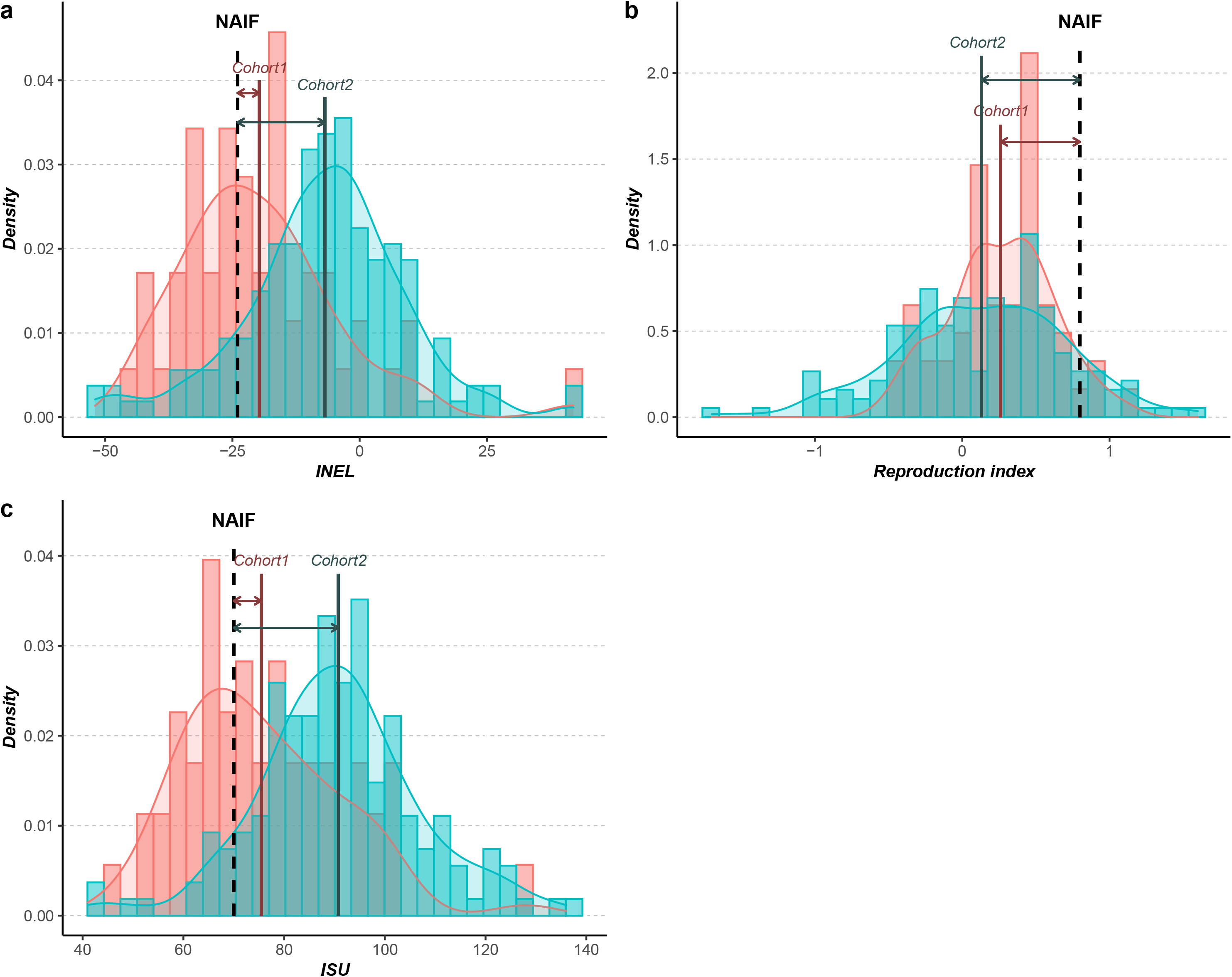
Distribution of INEL (a), reproduction index (b) and ISU (c) for Cohort 1 and Cohort 2. The 62 bulls in Cohort 1 are shown in pink, the 165 bulls in Cohort 2 are shown in blue, Naif is represented by the black dashed line, the mean of each cohort is represented by the solid line and the different gap between Naif and the mean of cohort by the arrows

The ISU, INEL and REPRO values of the 15 sons of Naif, from its second use, were 84.74 (sd=16.48), −11.40 (sd=17.86) and 0.67 (sd=0.33), respectively. The values of the 25 grandsons of Naif, from its second use, were 106.52 (sd=11.36), 6.88 (sd=12.93) and 0.27 (sd=0.27) for ISU, INEL and REPRO respectively. The ISU, INEL and REPRO values of the 155 bulls of the cohort 2 with no genetic link to Naif’s second use, were 91.14 (sd= 16.11), −6.38 (sd= 15.62) and 0.09 (sd=0.56), respectively. The values were 98.15 (sd=17.11), −1.64 (sd=15.82) and 0.08 (sd=0.48) for their 416 sons and were 107.82 (sd= 12.10), 6.12 (sd=11.91) and 0.06 (sd=0.36) for their 528 grandsons for ISU, INEL and REPRO respectively. These average values for the three indices and the relative position of Naif are shown on Figure 8. For the three indices at the generation 0, the values of Naif were significantly lower than the other bull means for ISU and INEL but significantly higher for REPRO (one sample t-test, t=16.33, df=154, p<0.05 for ISU, t=14.05, df=154, p<0.05 for INEL and t=-15.72, df=154, p<0.05 for REPRO). At generation 1, the means of the Naif lineage were significantly lower than the other bull lineages for ISU and INEL, and significantly higher for REPRO (F=8.93, df=1, p<0.05 for ISU, F=5.46, df=1, p<0.05 for INEL, F=22.42, df=1, p<0.05 for REPRO). At generation 2, the means between the two lineages were not significantly different for ISU and INEL (F=0.28, df=1, p=0.60 for ISU, F=0.10, df=1, p=0.76 for INEL), but the Naif lineage was significantly higher for REPRO (F=8.61, df=1, p<0.05 for REPRO). Moreover, we have computed the empirical within-family genetic variances of bulls’ GEBV for the three indices within each sire families present in the pedigree at generation 1. The ISU, INEL and REPRO variances between the male progeny of Naif (i.e. half-sib family) were 271.64, 319.11 and 0.11 respectively, while the average values for the other sires male half-sib families were 194.80, 190.72 and 0.21 respectively.

**Figure 8.**
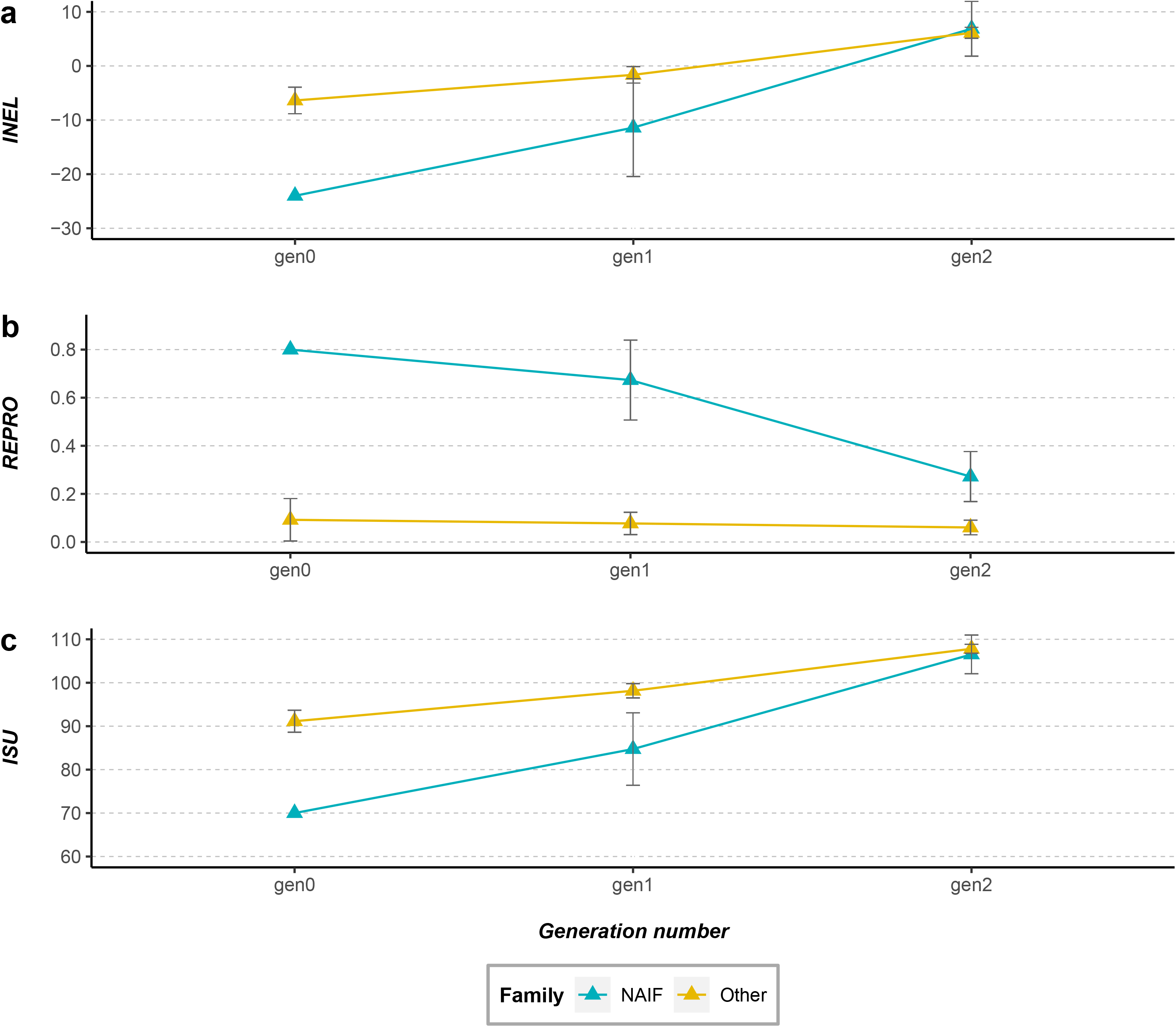
Average values of INEL (a), reproduction index (b) and ISU (c) at two generation after the second use of Naif. The average values of the offspring from the reuse of Naif’s frozen semen are represented in blue and those of the offspring from the other sire families are represented in yellow. Error bars correspond to confidence interval

### Females

The INEL and MILK values of the cohort 1a were −30.44 (sd=18.34) and −700.81 (sd=451.55) respectively. Average values for the cohort 1b were −30.19 (sd=15.74) and −625.84 (sd= 377.57) for INEL and MILK respectively. The INEL and MILK values of the cohort 2a were −7.00 (sd=16.36) and −188.49 (sd=404.63) respectively. Average values for the cohort 2b were 5.90 (sd=16.27) and 157.28 (sd=309.49) for INEL and MILK respectively. The distribution of these values for cohort 1a and cohort 2a and the relative position of the average values for the cohort 1b and cohort 2b are shown on Figure 9. The difference between the mean of cohort 1a and 1b for MILK and INEL, were −74.97 and −0.25, respectively, and −345.77 and −12.90, respectively, between the mean of cohort 2a and 2b. For INEL and MILK indexes, the means of cohort 1a and cohort 1b were not significantly different but the means of cohort 2a and cohort 2b were significantly different (F=978.2, df=3, p<0.05 for INEL and F= 768.6, df=3, p<0.05 for MILK).

**Figure 9.**
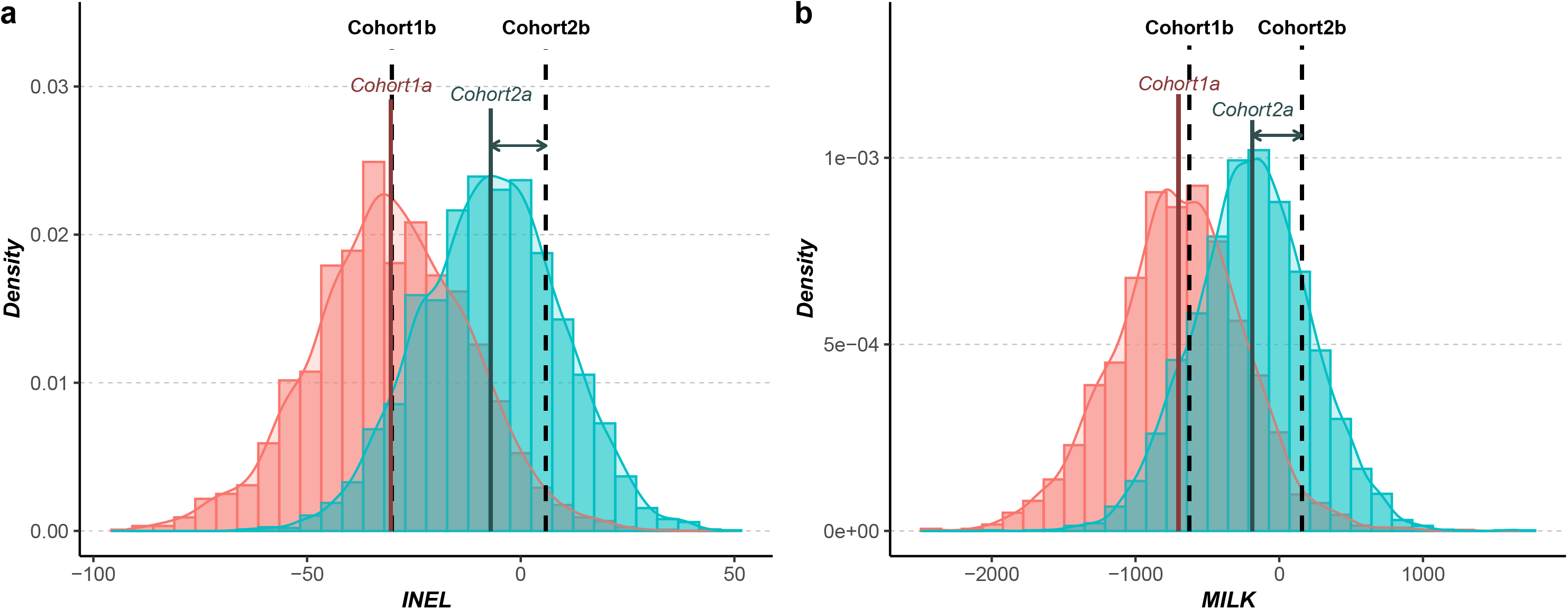
Distribution of INEL (a) and a milk index (b) for females mated with Cohort 1 and Cohort 2. The 2443 cows in Cohort 1a are shown in pink, the 4092 cows in Cohort 2a are shown in blue, the performance mean of the Cohort 1b and Cohort 2b are represented by the black dashed lines, the mean of each cohort is represented by the solid line and the gap between the different cohort means by the arrows

## Discussion

The AI bull called Naif (born in 1977), belonged to a disappeared family, namely, the progeny of the bull called Amiens. This bull was chosen to help to “purify” the breed, i.e. to contribute to decrease the proportion of Red Holstein breed within the Abondance breed. Naif had also the advantage of having a large stock of cryopreserved semen. He was therefore a good candidate to successfully bring back genetic diversity, especially genetic origin from the Amiens lineage. Naif procreated 45 offspring in the first use and, after a ten years period of inactivity, its semen was used once again to produce 33 offspring in the second use but in a shorter time period indicating a more intense use of its semen. The years from 2004 to 2007 were marked by an important production of individuals with Naif as a sire with a total of 30 descendants in those 4 years only. These two successive uses let to a large genetic contribution of Naif to the population. Its overall contribution increased in both periods of use with stagnation when not used, probably because its offspring was not used anymore to produces sire bulls since its lineage was abandoned. However, his contribution continued to increase after 2009 when he was no longer used as a sire, through the use of Naif’s progeny into the Abondance breeding scheme. Moreover, while the old contribution remained almost stable from 2004 to 2017, recent contributions increased over the end of this period. Indeed, the recent contribution of Naif became almost equivalent to the old contribution from 2016 onwards. From 2013 onwards, the recent contribution increased year by year due to the use of sexually mature offspring from the second use of Naif. Thus, the evolution of the recent contribution of Naif showed that descendants of contemporary use of Naif have not been excluded from the Abondance selection scheme.

The performance study revealed that Naif was an average individual during its first use. However, a discrepancy was observed when compared to a more recent cohort. Indeed, when reintroduced, Naif had much lower performances for production traits as observed for total merit index, ISU, and milking merit index, INEL, than the sires used during its second use. A gap of nearly 2 generations between cohorts 1 and 2 resulted in an increase of 15.29 points and 12.86 points for the ISU and INEL respectively. Such a lag in performance in a selected breed was expected and has been already highlighted in previous studies [14,15]. This difference is likely to be even greater under more intensive selection, resulting in larger genetic gain, like for mainstream dairy cattle breeds such as the Holstein breed. Thus, the larger the annual genetic gain, the more detrimental the effect of older individuals on the performances for the selected traits. Leroy *et al* [15] showed by simulations that an effective integration of cryopreserved sires with lower genetic values into breeding schemes would only be possible through an effort to conserve their progeny for future mating. In the case of Naif, we revealed that this gap in performances can be effectively absorbed in few generations through appropriate mating plans with performant females that could compensate this lag. Indeed, we showed that females that have been mated with Naif during its second use exhibited larger breeding values than females mated with other contemporary bulls. Moreover, within two generations, the difference in breeding values had already begun to be amortized. While Naif was 23.2% and 276.1% lower for ISU and INEL, respectively than the sires used in his second use, this gap was only 1.2% and 12.4% for ISU and INEL after two generations, respectively. However, a latency period is necessary in order to fill this performance gap, mainly due to both generation interval and the magnitude of this performance gap. In plant breeding, Allier *et al* [16] showed that collaborative diversity panels (i.e. genetic resources and elite lines) coupled with genomic prediction seemed relevant to identify and exploit genetic resources to enrich elite germplasm in maize. However, they showed that “bridging” and “pre-breeding” steps favors the insertion of genes from genetic resources by limiting the mismatch on the traits of interest for breeding [17]. Then the qualities brought back by Naif could be integrated into the breeding scheme mainly through its progeny. This was revealed by the continuously increasing genetic contribution of Naif to the Abondance breed after the end of its second use, and in particular its increasing contribution from 2013. Thus, as suggested by Eynard *et al* [14], we proved from field data that adding older bulls to a current breeding population undergoing selection was feasible without a too large impact on the genetic merit. In addition, the high heterozygosity of Naif could give access to a wide gametic variance (i.e. a large genetic variability of its gametes). Associated with a large number of offspring when it was re-used, the gametic variance allowed it to offer a rather large panel of diversity among its descendants that were hereafter selected in the breeding scheme. Indeed, the empirical genetic variances for the three indices (INEL, REPRO and ISU) of the sons of Naif after its reuse were all higher than the corresponding average within-family genetic variances of the other contemporary bulls. Other studies have also revealed that the use of parents who produced more variable gametes may provide a better response to selection by increasing the probability of reproducing a high-level genotypes [31,32].

However, in spite of the negative impact on performance traits that have been under constant selection, the use of Naif has increased qualities in other traits, in particular the ones that were not or weakly selected. In our study, the use of Naif brought performances for reproduction traits whose improvement was not necessarily as strong within the past breeding goals as for production traits. Indeed, while Naif was 767.1% ahead of the other sires used at the time of its second use for REPRO, after two generations, its progeny was still 350.2% ahead. The progress made on a production index like INEL, but also the advance kept on the REPRO index allowed to absorb the delay of Naif’s progeny on the ISU total merit index. This underlines the fact that traits such as fertility, calving ease or vitality at birth, which are strongly affected in many dairy breeds by the selection strongly focused toward production traits [33–35], can be improved using cryopreserved resources. The change in breeding goals is one of the case highlighted by Leroy *et al* [15] in their simulations that make old genetic resources valuable in a breeding scheme.

The study of the 2017 population, nearly two generations after the second use of Naif highlighted the success of the reintroduction of diversity within the breeding scheme. Inbreeding did not increase, a slight decrease in inbreeding between individuals resulting from recent Naif matings and the rest of the population was even observed. This significant difference in inbreeding can be explained by the choice of females that were mated with Naif. Indeed, the action of bringing out an old bull can favor inbreeding if mating were conducted with related individuals (i.e. its descent). This can be even stronger if the cryo-preserved individual has been heavily used in the past. Conversely, in the case of Naif use, we observed a decrease in inbreeding of offspring from the re-use of its semen. The present study case was probably favorable to this since the Amiens family to which Naif belongs to has almost disappeared from the breeding scheme, or at least was not used anymore to produce sires. More generally, the longer the time lapse between first and second use of sires, the less strong is the kinship, and thus the impact on the inbreeding of the population. In addition, Doekes *et al* [36] showed that recent inbreeding is more detrimental than the one due to more distant generations, since it leads to more inbreeding depression in many traits due to a shorter action of selection against inbreeding load (i.e. purging).

Finally, multivariate analyses revealed genetic uniqueness of Naif. When first used, Naif was representative of its contemporary sires. However, when used for the second time, Naif appeared to be an original individual relative to the other active sires. The BCA computed two generations after the last use of Naif reveals that all individuals with a recent link to Naif are distinguishable from other individuals, thus the genetic originality of Naif has been transmitted to Naif’s descent. While the initial motivation was a return to “pure” breed, the use of an old cryopreserved individual was thus able to bring part of the lost genetic specificity of a lost family back into current population and thus reversely increased genetic variability.

## Conclusions

This study is one of the first to document a real case of the long-term impact of the use of cryopreserved reproductive material in a domestic animal breeding scheme, both in terms of performances and genetic diversity. We showed that despite a lag in yield traits, the use of appropriate mating plans can fill this gap in a few generations. Furthermore, these disadvantages can be balanced by advantages on other traits for which breeding goals have changed since the conservation of genetic resources, for instance in the case of functional traits. In terms of genetic diversity, the use of cryopreserved material from ancient individuals can be beneficial by providing genetic originality to the current population that could have been lost over time. Additionally, the effectiveness of the use of old germplasm for genetic improvement also depends on the gametic variance of the reintroduced material that should be as high as possible. It should be checked at least through the heterozygosity or, when feasible, through the haplotype diversity of donors. In terms of genetic diversity, the use of old cryopreserved individuals was beneficial by providing genetic originality to the current population that can be lost over time. Moreover, inbreeding rate can be controlled using planned matings with non or weakly related individuals. Thus, particular attention should be paid when re-using a sire that has been massively used in the past. Reversely, the use of extinct lineages or very original individuals should have a higher positive impact on genetic diversity. In this respect, molecular data obtained from routine genotyping can be used to characterize the level of genetic diversity that could be brought to the current population by sires from a gene bank. Indeed, genomic selection allows for the evaluations of former individuals on the same scale than the one of the current populations, even for traits that were not selected in the past. Considering that cryopreserved genetic resources are seldom used nowadays in breeding programmes, we provide here some clues to explain how successful the use of cryopreserved material can be. Further studies are still needed to tailor the recommendations for using cryopreserved resources to the different objectives and expectations of each species and breed (e.g. management of a genetic defect, improvement of disease resistance, diversity reintroduction…). Finally, our approach could be easily transposed to wild populations for which genetic diversity reintroduction from zoos or reserves is a major concern.

## Authors’ contributions

GR et GL conceived the project. AJ performed the analysis and drafted the manuscript. GR supervised the analyses. AJ, MTB and GR interpreted the results. AJ, GR, GL, XR, EV and MTB contributed to the writing of the manuscript. All authors read and approved the final manuscript.

## Acknowledgements

The authors would like to acknowledge the organization for the selection of alpine breeds (OSRAR) for their comments and suggestions on this study.

## Additional files

**Additional file 1 Figure S1**

Format: jpeg

Title: Cows from Abondance breed ruminating on high altitude pastures in the Chablais mountains, © Étienne Verrier (August 2022)

**Additional file 2 Figure S2**

Format: pdf

Title: Classification of different genetical links with the Naif bull for the cohort 2017

Description: Red: individuals with no genetic family link to Naif (0_LWN), Purple: individuals with a recent genetic link with Naif through one of their parents (1_LWN_R), Green: individuals with an old link with Naif through one of their parents (1_LWN_O), Orange: individuals related to the first use of Naif by both parent (2_LWN_O), Magenta: individuals related to the recent use of Naif by both parent (2_LWN_R), Blue: individuals with one old and one recent genetic link with Naif (2_LWN_OR)

**Additional file 3 Figure S3**

Format: pdf

Title: Principal Component Analysis of genotyping data for Cohort 2

Description: Green: individuals with no genetic link to Naif (noLWN) - Red: individuals with a genetic link to Naif (LWN) - Naif is represented by the blue dot

