## Supplementary figures and images for "Reintroducing genetic diversity in populations from cryopreserved material: The case of Abondance, a French local dairy cattle breed"

### Figure S1

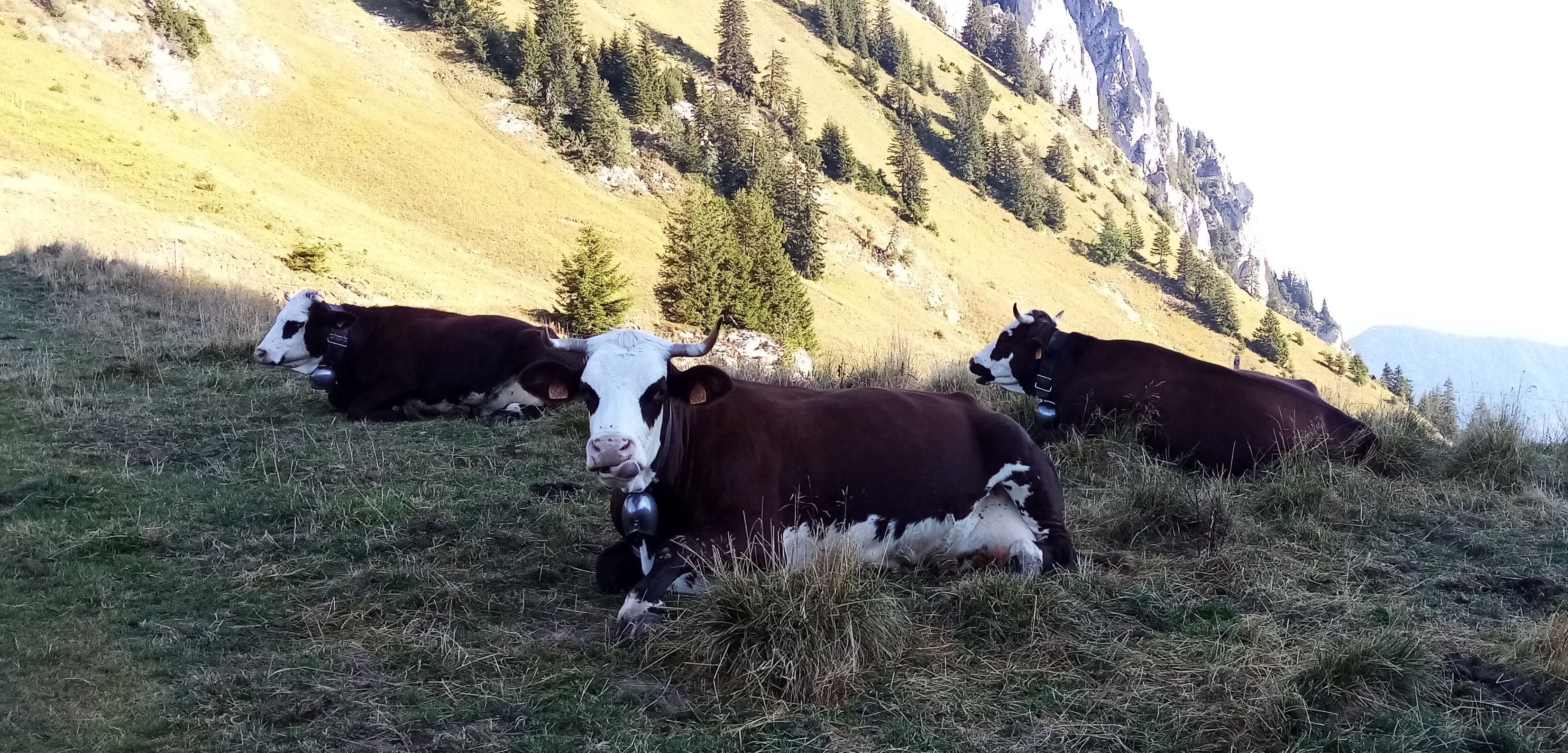

### Figure S2

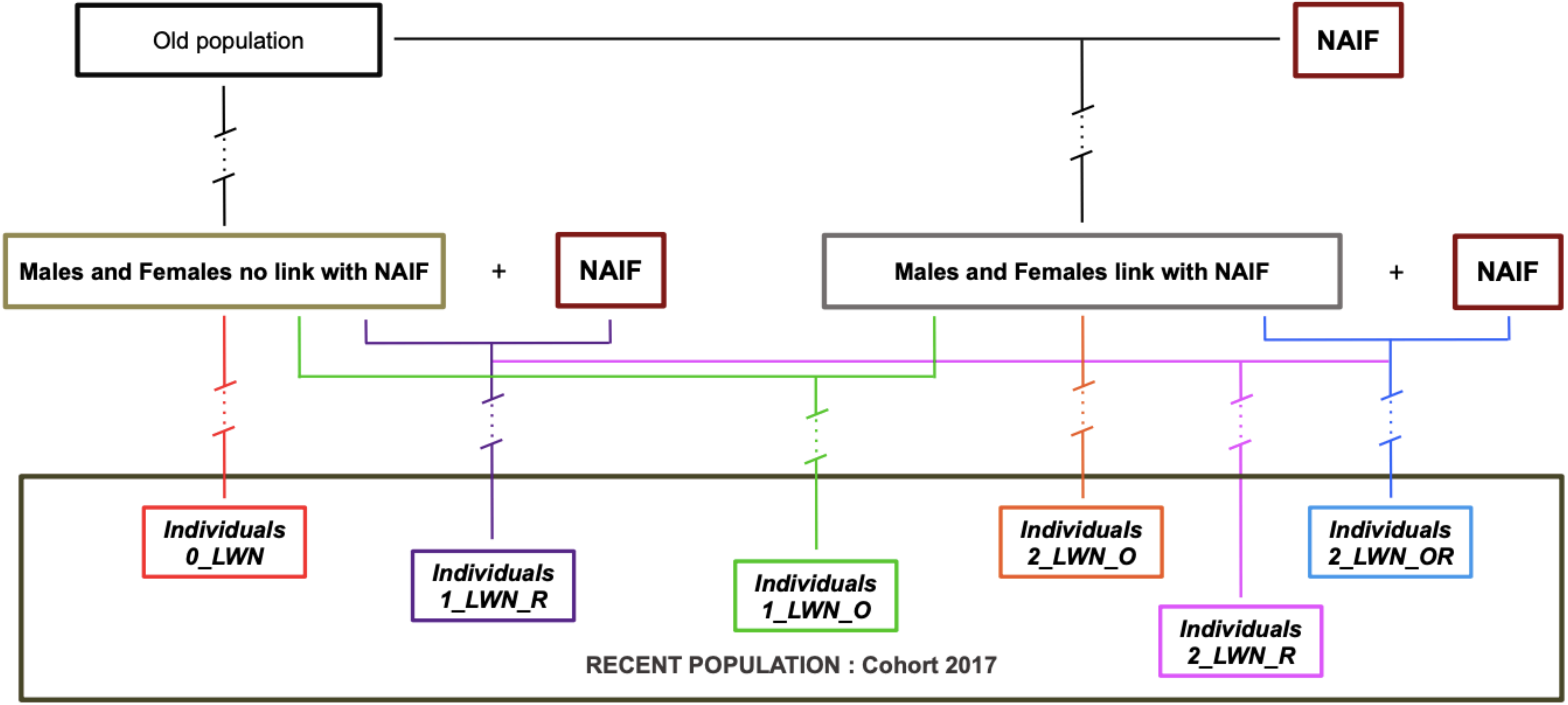

### Figure S3

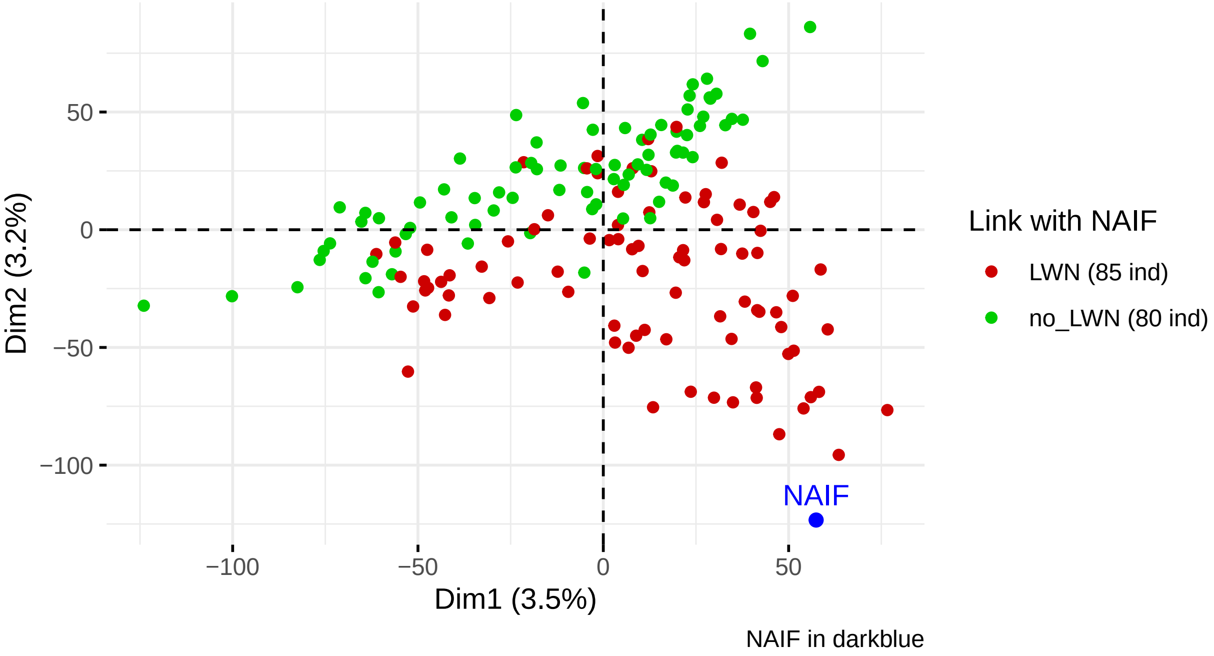
